# Deeply-sequenced metagenomes and over 1000 draft genomes from the epipelagic to bathypelagic Northeast Pacific Ocean

**DOI:** 10.64898/2026.07.18.739368

**Authors:** Alexander L. Jaffe, Rebecca S.R. Salcedo, Jasmine J. Lewitton, Anne E. Dekas

## Abstract

The deep ocean water column (>200 meters depth) hosts diverse and active microbial communities that play critical roles in Earth’s biogeochemical cycles. However, this habitat has been massively undersampled compared to shallower depths where sunlight penetrates, as well as benthic features like hydrothermal vents and seeps. In particular, few existing deep-sea datasets provide adequate spatial resolution to study the impact of physical and geochemical gradients on marine microbial communities over both horizontal and vertical spatial scales. Here, we present a deeply-sequenced, spatially-resolved set of 28 water column metagenomes from a ∼300 km horizontal transect off the coast of central California, spanning 50 to 4000 meters depth. We estimate that the deepest portion of the resulting ∼1.5 terabases of sequencing data comprises nearly 9% of all shotgun metagenomic sequencing currently available from 1000 meters depth or below. Additionally, we generate over one thousand draft genomes describing the bacteria and archaea present in our bulk metagenomic data. These genomes frequently account for the majority of reads sequenced, and show discernable patterns of abundance with depth, underscoring the presence of distinct microbial cohorts inhabiting different ocean compartments. Additionally, the depth of sequencing performed (mean of ∼51.9 gigabases per sample) enables resolution of rare community members that are, on average, more phylogenetically novel than their abundant counterparts, forming an important contribution to genomic catalogues for this habitat. We anticipate that the combined datasets will enable diverse research questions concerning the adaptation, functional potential, and distribution of deep sea microbes as well as their genes and proteins.

## Background & Summary

Since the advent of the field in the late 1990s and early 2000s^1–3^, community DNA sequencing has helped to reveal the tremendous diversity and complexity of marine microbiomes as well as their relationships to global physical, chemical, and climate processes. Both large scale oceanographic surveys (e.g. Tara Oceans, Malaspina, GEOTRACES)^4–6^ as well as smaller, individual sampling efforts have contributed to an ever-growing corpus of genomic data that describes the bacteria, archaea, viruses, and smaller eukaryotes comprising ocean microbial communities^7–9^, and have allowed countless insights into their ecological and biogeochemical roles. However, due primarily to difficulty of sampling, as well as the lower density of cells, microbial communities in the deep sea (>200 meters depth) are highly undersampled compared to the surface ocean^10–13^, where sunlight penetrates and supports orders of magnitude higher degrees of biomass^14^. Similarly, the deep water column is undersampled relative to marine sediments, hydrothermal vents, seeps, and ridges due to their unique geochemistry, which is often able to support high cell densities compared to surrounding waters^15^.

Existing datasets suggest that microbial communities in the vast, deep pelagic water column are diverse, active, and play significant roles in elemental cycling, but leave many biological questions unanswered. This motivated us to study microbial communities in the deep ocean off the coast of central California in relation to the physico- and geochemical features of the region (Fig. 1ab). In previous work, we have drawn upon data from this cruise to study the diversity, activity, and genetic features of a subset of the ocean microbiome. Specifically, we have previously examined general patterns of microbial activity across the transect^16^, primarily by single-cell microbial activity assays, and have also applied metagenomics to identify specific lineages assimilating urea^17^. Additionally, we have also leveraged a subset of the data reported here to investigate the diversity, abundance, and metabolism of organisms capable of autotrophy across the transect, including (but not limited to) autotrophy mediated by organisms encoding the enzyme rubisco^18,19^.

**Figure 1.**
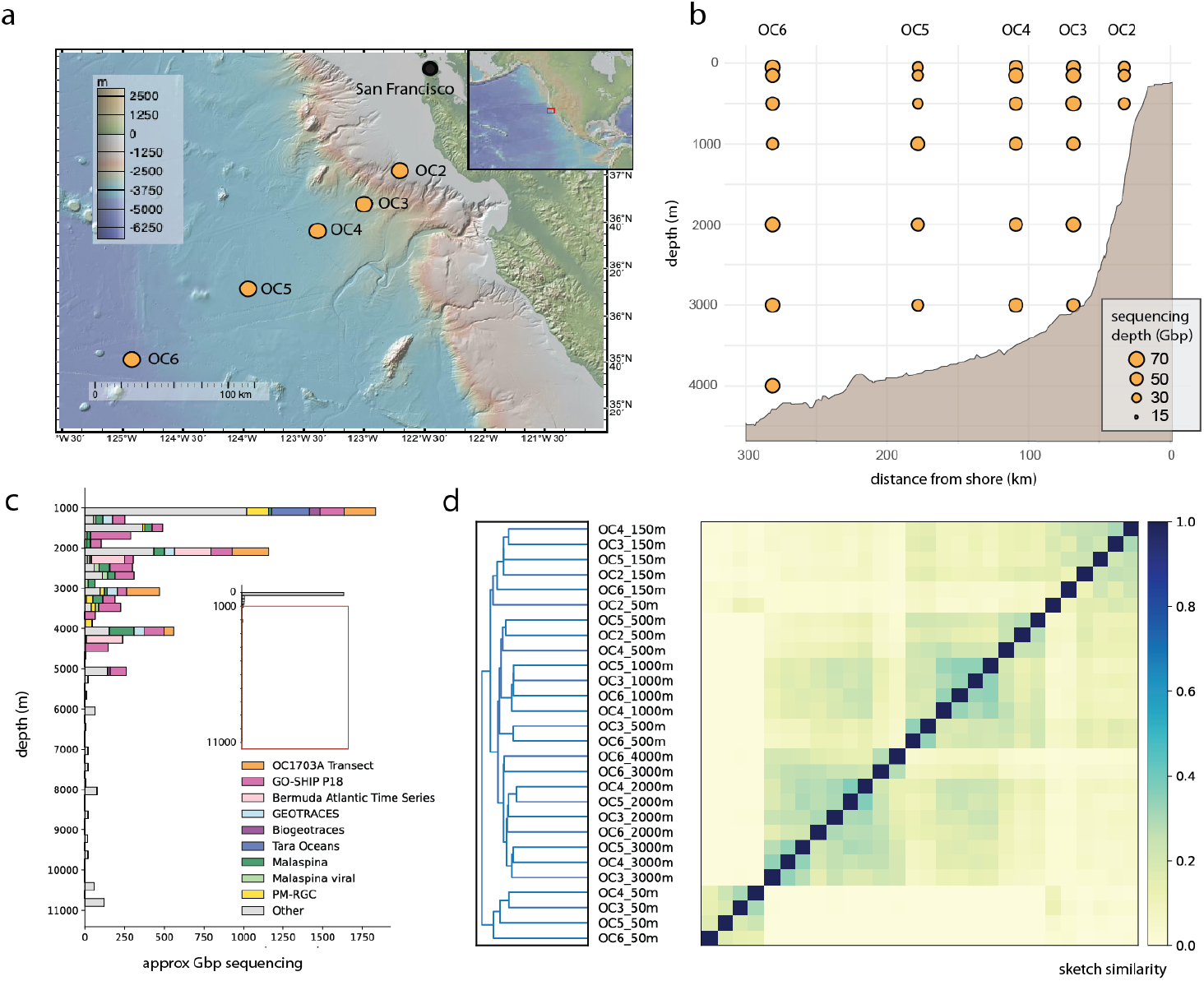
Characteristics of the OC1703A Transect metagenomes. **a)** Cruise track and regional bathymetry produced using GeoMap App (v3.7.5). **b)** Spatial distribution of samples collected for metagenomic sequencing, as well as sequencing depth per sample in gigabasepairs (Gbp). **c)** Metagenomic sequencing effort below 1000 meters depth represented by this dataset relative to those from the public record (Methods); inset depicts full depth range from the surface to ∼11,000 meters. **d)** Patterns of metagenome similarity across the transect based on k-mer containment (Methods).

The lineages so far analyzed from the OC1703A Transect dataset represent a minority of its microbial community and, further, a small fraction of the total assembled metagenomic sequence. Here, we reiterate the 28 metagenomes sequenced as a part of this research effort (Table S1) and newly contribute the corresponding metagenomic assemblies as well as 759 manually refined draft genomes as resources for the scientific community. Based on analyses of comparable public data available through the National Center for Biotechnology Information (NCBI), we estimate that the deepest portion of our sequenced samples comprises approximately 9% of shotgun metagenomic sequencing data currently publicly available for untreated marine samples from 1000 meters or below (Fig. 1c, Table S2). In addition, we present a subset of contigs from the assembled sequence that can serve as a starting point for analyses of the viral community within our samples. We expect that the combined dataset, alongside the previously published activity and geochemical data^16,17^, will make valuable contributions to future studies of the ecology and evolution of diverse microbial lineages as well as their genes and proteins far beyond the initial scope of our work.

## Methods

### Sample collection, extraction, and sequencing

Seawater samples were collected off the central California coast from the *R/V Oceanus* from the 12th to the 23rd of March 2017. At each of 6 sites along a ∼300 km transect (OC2, OC3, etc), 12 L Niskin bottles were deployed to collect discrete seawater samples at 50, 150, 500, 1000, 2000, 3000, and 4000 meters depth, as total ocean depth permitted (Fig. 1ab). Seawater (5–25 l depending on depth) was filtered using a peristaltic pump (Cole Parmer, USA) onto 0.2 µm Sterivex units (Millipore, Germany). Filters were harvested from the units, placed in 2 mL nalgene cryovials, and flash frozen in liquid nitrogen before storage at −80°C.

Back onshore, filters were removed from their casings and DNA was extracted using the AllPrep DNA/RNA kit (Qiagen, Valencia, CA, USA). After the elution step, DNA was quantified using a Qubit fluorometer with the high-sensitivity, dsDNA reagent/standards (Invitrogen, Waltham, Massachusetts) and purity was assessed using a Nanodrop spectrophotometer (Thermo Fisher, Waltham, Massachusetts). Input DNA was normalized to 8 ng per sample to account for low yield and subsequently prepared using the Takara ThruPLEX kit (Takara Biosciences USA, Mountain View, CA) with 10 PCR cycles. A PippinHT (Sage Science, Beverly, MA) was used for size selection. Size-selected libraries were ultimately pooled and sequenced on a NovaSeq 6000 S4 platform (150 bp paired-end reads) at the UC Davis DNA Sequencing Facility.

### Sequencing quality control and metagenomic assembly

Raw sequencing reads were trimmed using BBDuk (https://bbmap.org) using the following parameters: *ktrim=r k=23 mink=11 hdist=1 tbo qtrim=r trimq=25 minlen=20*. Trimmed reads were then assembled individually using MEGAHIT^20^ (v1.2.9) with the *meta-sensitive* preset, and filtered to resulting contigs attaining ≥1000 bp in length. To visualize high-level sequence relationships between metagenomes, sourmash^21^ (v3.5.1) sketches were computed for each assembly using default parameters, and compared using a k-mer size of 31 (Fig. 1d).

### Contextualizing the OC1703A Transect sequencing effort

To compare our sequencing effort to comparable datasets in NCBI, we constructed a compound search query to the Sequence Read Archive (SRA):

*marine metagenome”[Organism] OR “seawater metagenome”[Organism] NOT benthic NOT sediment NOT symbiont NOT “coral holobiont” NOT amplicon[Strategy] NOT “rna seq”[Strategy] AND metagenomic[Source]*

This query was designed to exclude large BioProjects containing host- or sediment-associated marine microbiomes in favor of those from the pelagic water column. Results were outputted in the SRA ‘RunTable’ format and read into a Jupyter Notebook with a Python (v.3.6.1) kernel. Where already existing, depth metadata were cleaned/reformatted using custom code (see Code Availability Statement). Depth metadata were manually curated for SRA accessions that appeared to originate from depths at or below 1000 meters or for BioProjects where sequencing runs cumulatively exceeded 1 gigabasepair of sequencing. Given the large number of accessions in this category, we prioritized those BioProjects that contained the terms “deep-sea”, “deep”, “mesopelagic”, “bathypelagic”, “abyssopelagic”, “hadal”, “hadopelagic” in their names, descriptions, or titles.

To focus on metagenomes derived directly from *in situ* seawater samples, we removed any that appeared to have been treated experimentally in any way, including incubations and amendments. Total sequencing effort was plotted as a function of depth and annotated with expedition/cruise of origin in select cases. Due to the inconsistent population of metadata fields within the SRA, particularly for newer projects, we acknowledge that this approach is non-exhaustive and relevant runs may be omitted due to incomplete metadata. Thus, we report our statistics as conservative maxima.

### Metagenomic binning and manual bin refinement

We first mapped trimmed reads to assembled sequences for a subset of sample pairs determined by computing overall assembly similarity using sourmash. To reduce the computational burden of the mapping step, only sample pairs attaining ∼10% similarity or greater were processed using bowtie2^22^ (v2.5.4), specifically:

- Assemblies from 50 meters depth, being highly distinct, were only used as alignment indices for reads from 50 and 150 meters depth
- Assemblies from 150 meters were used as alignment indices for reads from 50, 150, 500, and 1000 meters depth
- Assemblies from 500 and 1000 meters were used as alignment indices for reads from 150 meters and below
- Assemblies from 2000, 3000, and 4000 meters were used as alignment indices for reads from 500 meters and below

Alignment results were converted to bam format using *samtools*^*23*^ (v1.16.1) and were ingested by metabat2^24^ (v2.12.1) for differential coverage binning.

Initial bins were evaluated for quality using CheckM^25^ (v1.1.10) and de-replicated using dRep^26^ (v3.4.0, *-sa 0*.*95*). Bins that met initial quality thresholds (≥50% completeness and ≤25% redundancy) and were selected as cluster representatives at 95% ANI were manually refined using the anvi-refine function from Anvi’o^27^ (v7). In this process, the results of which are reported in Table S3, contigs belonging to a bin were clustered by their sequence characteristics (GC content, tetranucleotide frequency, coverage across samples) and were visualized as a dendrogram. Contigs were removed using the visual interface if they deviated from typical GC content or coverage profiles describing the plurality of contigs. Generally, a conservative approach was taken, such that sets of contigs that decreased overall bin completeness by more than 15% were not removed. Full custom code required to perform manual curation is linked in the Code Availability section below.

After manual curation, CheckM was used to re-assess quality metrics and to filter down refined genomes to those attaining ≥50% completeness and ≤10% redundancy. We also subjected genomes to a second round of dereplication using dRep (95% ANI) to yield a final, representative set of 1,004 genomes. Representative genomes were additionally profiled using GUNC^28^ (v1.0.6), which examines taxonomic profiles across contigs to identify ‘cryptic’ contamination not necessarily determined by marker gene approaches. Differences in quality status before and after manual curation were visualized using the seaborn library in Python (v0.11.2). Final taxonomic assignments were inferred using GTDB-Tk (v2.4.1) *classify_wf* workflow with database release 226^29^. rRNA and tRNA gene numbers were computed using *barrnap* (v0.9) (https://github.com/tseemann/barrnap) and tRNAscan-SE^30^ (v2.0.12), respectively. For tRNAs, multi-copy genes were disregarded, and only genes associated with the 20 canonical amino acids were considered. According to MIMAG criteria^31^, approximately 3.4% of representative genomes were considered high-quality, while the remaining representative genomes were considered medium-quality (Table S4).

### Evaluation of assembly and binning efficacy

To assess the efficacy of our assembly and binning processes, several analyses were undertaken. First, we mapped all trimmed reads against the representative genome set as well as each set of assembled contigs using bowtie2. We then computed the percentage of trimmed reads recruited by the genomes and assemblies, respectively, using CoverM^32^ (v0.6.1) with loose mapping parameters (*--min-covered-fraction 0 --min-read-percent-identity 0*). Estimates of the ‘prokaryotic fraction’ – the percentage of reads estimated to derive from bacteria and archaea – were made using SingleM^33^ (v0.20.3) and were used to adjust the read mapping rate for genomes using the following formula:

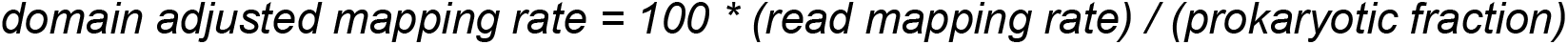

Bulk community composition was inferred by applying the SingleM *pipe* function to the trimmed reads (*--assignment-method diamond*) using a custom metapackage amended with the refined genome set (*supplement* function). The *pipe* function was also applied to the assembled contigs and refined genomes, and OTU tables from all three data types were passed to the *appraise* function (*--imperfect*) to generate plots depicting the retention of OTUs through the assembly and binning process. Full code describing this process is linked in the Code Availability section.

### Analysis of divergence and distribution for recovered genomes

To examine patterns of diversity and abundance across the transect, reads were re-mapped to the representative genome set using bowtie2 and filtered stringently using CoverM (*--min-read-percent-identity 0*.*95 --min-covered-fraction* 0.50). Relative coverage was calculated by dividing mean coverage values for each genome by the summed coverage of all genomes detected (i.e. recruiting reads at or above the identity and breadth thresholds) within a sample. To assess the divergence of representative genomes from known references, a fastAAI (https://github.com/cruizperez/FastAAI) database was constructed using genomes from GTDB release 226. Representative genomes were then compared to the database, and the best match per genome was selected. Relative coverage and AAI values were then visualized as a function of sample, depth, and percentile abundance across all transect samples using Python.

### Viral identification and analysis

We began by screening our assemblies for viral sequences using geNomad^34^ (v1.8.1) and VIBRANT^35^ (v1.2.1). To maximize the recovery of potential viral sequences, output sequences from both tools were merged. We next clustered the merged sequences using vclust^36^ (v1.3.1) to remove duplicates and near-identical contigs, producing a set of representative viral contigs for each sample. The quality of these representatives was evaluated using CheckV^37^ (v0.8.1). Finally, we quantified viral abundance by mapping metagenomic reads from each sample back to its representative set of contigs using loose mapping parameters (*--min-read-percent-identity 0 --min-covered-fraction 0*). Characteristics of putative viral genomes – including sequence features, initial taxonomy, and quantification metrics – are provided in Table S5.

## Data Records

Cruise and environmental metadata are available in Table S1 as well as the NSF Biological and Chemical Oceanography Data Management Office (BCO-DMO) (#710001). Metagenomic sequencing reads and refined genomes in FASTA format are available through the NCBI under BioProject accession PRJNA1054206. Individual accessions for reads and draft genomes are available in Tables S1 and S4, respectively. For brevity, column headings are defined in the ‘Column Headings’ tab of the Supplementary Tables file. FASTA-formatted draft genomes and assembled contigs are also available through Zenodo.

## Data Overview

To further contextualize the full set of draft genomes presented here, we profiled their abundance and distribution across the transect samples (Fig. 2ab, Table S4). Emerging from these data were clear genome cohorts with similar vertical distributions through the euphotic, mesopelagic, and bathypelagic (as well as combinations thereof), consistent with the notion of depth as a driver of microbial diversity in marine systems^38–40^ (Fig 2a). Select individual taxa include apparent mesopelagic specialists (Fig. 2b, top panel) as well as bathypelagic specialists (Fig. 2b, bottom panel). The presence of distinct spatial ecotypes suggests that our sampling efforts targeted distinct, endemic microbial communities across depths, and thus will allow future work probing the genomic bases of adaptation to various physico- and geo-chemical gradients in the deep sea, including hydrostatic pressure.

**Figure 2.**
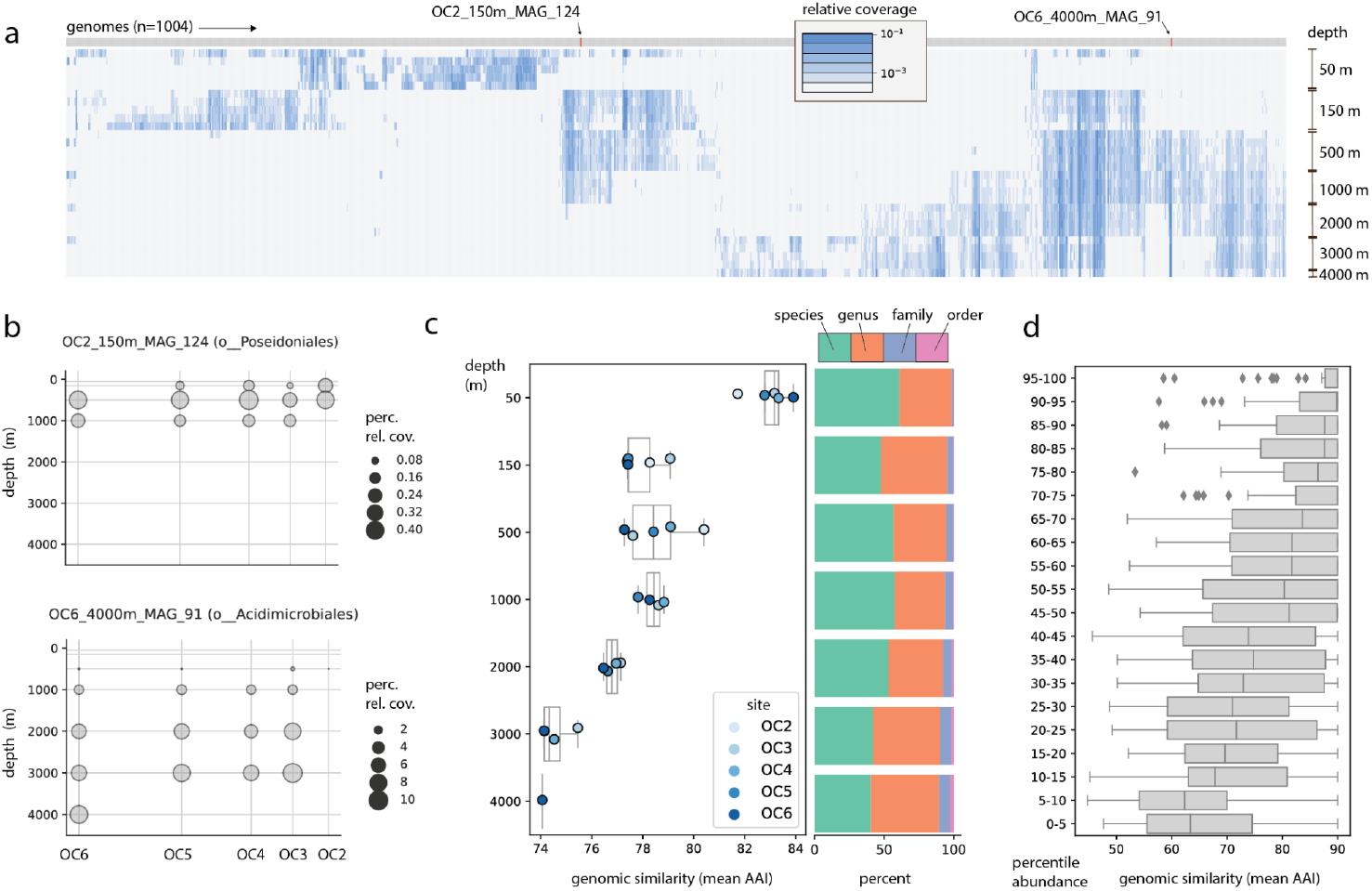
Patterns of abundance and similarity within the OC1703A Transect genome set. **ab)** Spatial distribution of draft genomes across transect sites/depths. Abundance values are expressed either as the **(a)** log-transformed proportion or (**b)** percentage of total sequencing coverage of all genomes detected within a sample. In **(a)**, each row represents a metagenomic sample (n=28) **c)** Mean genomic similarity of detected genomes at each site/depth relative to GTDB references. The stacked bar plot at right depicts the lowest taxonomic level at which detected genomes could be classified. **d)** Genomic similarity to GTDB references as a function of percentile mean abundance across all transect samples.

Additionally, we quantified the genomic divergence of each draft genome by comparing their encoded proteins to those from reference genomes (Methods). In general, we observed many genomes (∼54.3%) that could only be classified at the genus level or above, suggesting significant genomic novelty in the deep sea. Intriguingly, mean genomic similarity decreased with depth, reaching a minimum of ∼74.1% AAI at 4000 meters, consistent with the overall trend of undersampling apparent below 1000 meters (Fig. 1c). Finally, we ranked genomes by their mean abundance across all transect samples, observing that median similarity to reference sequences decreased with abundance percentile (Fig. 2d); or, in other words, that rarer microbes were more poorly represented in reference databases. This finding may be explained by the fact that typical sequencing allocations in the marine realm do not permit broad genomic resolution of rare microbiome members. Together, our results indicate that both the physical depth of seawater sampling as well as the depth of sequencing performed allowed us to recover quality genomic information for novel marine microbes, data which will undoubtedly benefit future studies of genomic diversity and evolution in various lineages.

## Technical Validation

Several steps were undertaken to ensure the quality of the genomic data produced. First, DNA extracts were quantified and measured for purity, ensuring that adequate material was put forth in the library preparation step. Libraries were re-quantified before pooling and sequencing. Once raw sequence information was generated, we visually inspected quality scores of both forward and reverse reads to ensure that most reads met a minimum average quality score of Q20 and that per-base quality did not decline beyond expectation into the read interior. Reads failing to meet quality standards were filtered out computationally (∼0.64 - 1.37% of reads, Table S1).

Once reads had been assembled and resulting contigs binned, we undertook a manual curation process to reduce the erroneous inclusion of genetic fragments within genomes. These ‘misbinned’ contigs can be highly problematic for downstream analyses, including (but not limited to) taxonomic assignment and metabolic reconstruction^41^. As described in the Methods section, contigs assigned to each bin were clustered by their coverage and sequence features and visualized using the Anvi’o web interface (example depicted in Fig. 3a). Contigs with abnormal coverage profiles and/or GC content were then removed, resulting in a refined set of draft genomes with improved redundancy metrics compared to their original versions (Fig. 3bc). While the mean improvement to genome redundancy metrics was modest (∼1.29% mean decrease in CheckM redundancy, ∼11.55% mean decrease in GUNC CSS), we observed that ∼72.47% of original bins were improved in one or both of these metrics after the process of refinement (Table S3). Notably, about 1.42% of bins were drastically improved (≥10% decrease in redundancy), and would have been discarded in their original state with the quality thresholds employed here (Fig. 3bc, Table S3). However, we did also observe small mean decreases to bin completeness during the process of curation (mean ∼1.86%), highlighting tradeoffs inherent to many current curation implementations (Fig. 3bc, Table S3).

**Figure 3.**
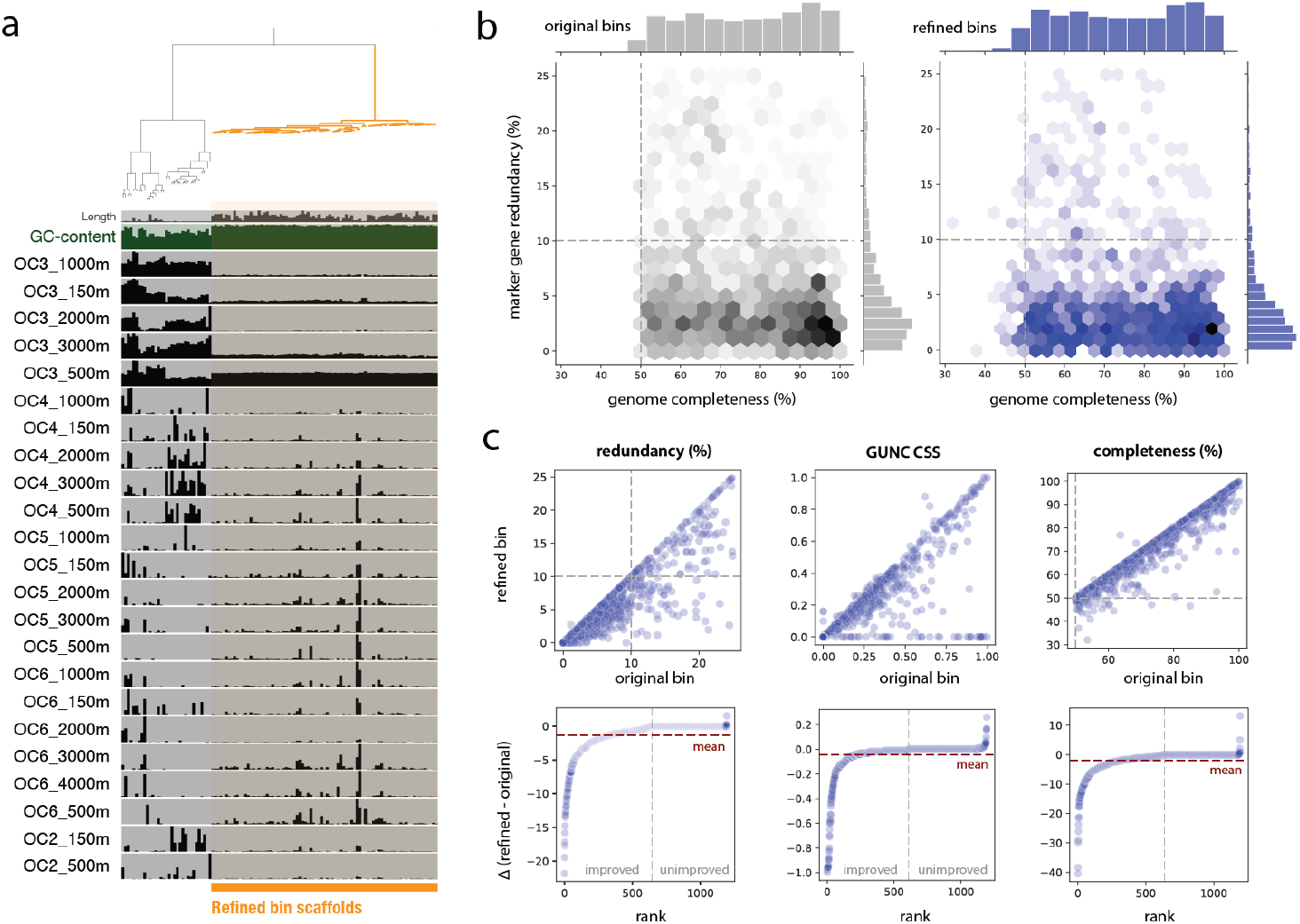
Genome refinement process and outcomes. **a)** example refinement process for a single original, unrefined bin, depicting the manual selection of a subset of binned contigs that were retained in the final, refined version. Shaded bars represent normalized coverage of each contig in a given sample. **b)** Quality metrics for the original and refined set of bins. **c)** Changes in CheckM completeness and redundancy, as well as GUNC Clade Separation Score (CSS), between the original and refined bins.

An important assessment of any metagenomic reconstruction effort is the degree to which it is representative of the microbial community originally sampled. To assess the community composition before any information loss during the assembly and binning steps, we profiled the trimmed metagenomic reads using a combination of conserved marker genes (Methods) (Fig. 4a, Table S6). We observed fairly stable populations of marine archaea and bacteria that were consistent with our previous investigations of the sample samples with 16S rRNA amplicon sequencing^16^. As suggested by our overall k-mer comparisons (Fig. 1d), 50 meter samples were generally distinct, with the exception of the sample taken nearest to shore (OC2_50m), where we hypothesize that upwelling transports microbial communities from deeper waters^42^. Based on read mapping, the majority of reads (median of 60.1%) were represented by the assembled sequence (Fig. 4b, Table S1). Similarly, a median of ∼51.2% of reads were represented by the refined genomes, once adjustments were made for non–prokaryotic sequence (median of ∼36.7% of reads without this adjustment)(Fig. 4b, Table S1).

**Figure 4.**
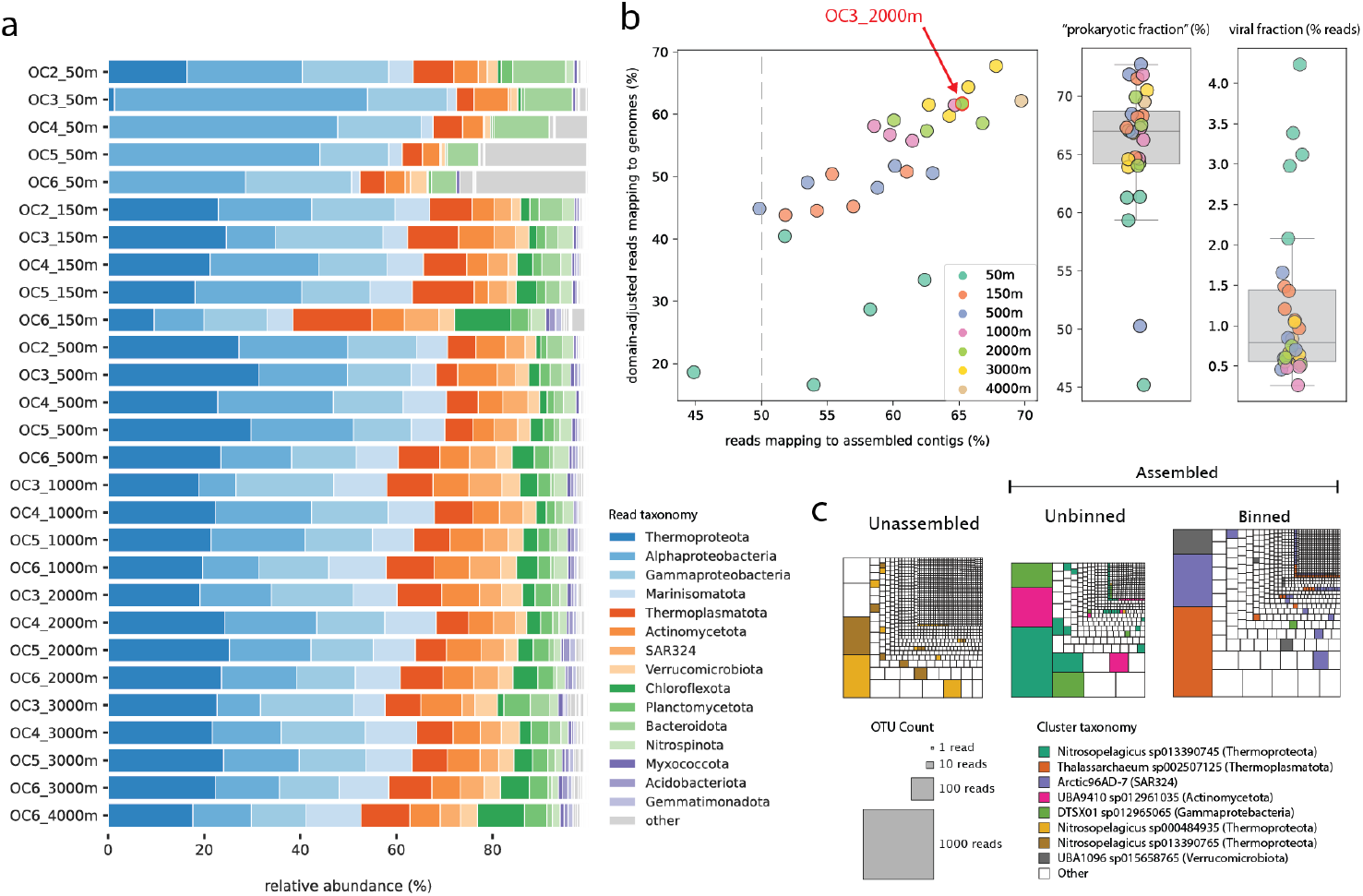
Genomic reconstructions of the Northeast Pacific microbiome. **a)** Community composition based on single marker genes extracted from reads. Only the top 15 lineages (phylum, or class in the case of Proteobacteria) are shown. **b)** Binning and assembly efficacy across transect samples, alongside estimates for the percentage of reads deriving from bacteria/archaea (prokaryotic fraction) and viruses. The highlighted sample (OC3_2000m) was selected for detailed analysis of lineage recovery/loss through the assembly and binning steps in panel **c)**.

To complement our estimates of the prokaryotic read fraction via SingleM, we also identified potential viral scaffolds and quantified their abundance using a custom bioinformatic workflow (Methods, Code Availability). We observed that these potential viral scaffolds recruited a median of 0.79% of trimmed metagenomic reads across samples, with substantially higher values observed in the shallowest samples at 50 meters (Fig. 4b, Table S5). We provide the full set of potential viral scaffolds, alongside their metadata, for future analyses of the viral community in this transect (Table S5, Data Availability Statement).

Finally, we tracked the representation of major taxa present in the reads through the assembly and binning process. While members of all major taxonomic groups were present throughout, we observed that certain lineages (e.g. abundant ammonia-oxidizing archaea from the genus *Nitrosopelagicus* or Alphaproteobacteria from the family Pelagibacteraceae, which includes SAR11) were often underrepresented in assembled and binned sequence, potentially due to high strain heterogeneity in these populations (Fig. 4c, Table S7)^43,44^. Other lineages like the SAR324 and *Thalassarchaeum* were more consistently assembled and binned. Table S7 provides a quantitative assessment of assembly/binning success for all lineages detected in the metagenomic reads, informing future re-assembly or binning efforts with alternative algorithms.

## Usage Notes

Despite our manual curation of over one thousand genomes, we cannot entirely rule out the possibility that some misbinned contigs are still present in the refined genome set. Given this, caution should be taken when performing analyses of genetic presence/absence (here or in any metagenomics dataset), particularly when small numbers of genomes or genes are considered, or contigs of interest are relatively short in length. In these cases, we encourage users to perform additional curation to ensure that genes of interest derive from the expected taxa. Additionally, as discussed above in the Technical Validation section, we acknowledge that not all microbial lineages identified in the read set could be captured in medium-to-high quality draft genomes, likely impacting any assessment of community composition based solely on assembled sequence. However, we provide a quantitative assessment of representation at each analysis step that can both guide targeted genome recovery and serve as a citable reference for specific taxa that are frequently underrepresented in marine metagenomes (Table S7).

## Data Availability

Trimmed metagenomic reads and draft genomes are available through NCBI at BioProject number PRJNA1054206. For redundancy and ease of access, FASTA-formatted draft genomes and assembled contigs are also available through Zenodo.

## Code Availability

Custom code is available via GitHub: https://github.com/alexanderjaffe/northeast-pacific.

## Author Contributions

A.E.D designed the sampling plan and collected samples. A.L.J. and A.E.D. designed the study. A.L.J., R.S.R.S., and J.J.L. contributed to genome curation, bioinformatic analysis, and the creation of figures. A.L.J. wrote the manuscript with contributions from all authors.

## Acknowledgements

We thank Alma Parada, Julian Fortney, and Nestor Arandia-Gorostidi for collecting and/or processing samples used for the metagenomic analysis. We thank the captain, crew, and science party of the *R/V Oceanus* OC1703A expedition. Finally, we acknowledge the Stanford Research Computing Center for informatic support as well as the Stanford Geomicrobiology Shared Laboratories Core Facility (RRID:SCR_025000).

